# *Listeria monocytogenes* infection in intestinal epithelial Caco-2 cells with exposure to progesterone and estradiol-17beta in a gestational infection model

**DOI:** 10.1101/2023.07.21.550068

**Authors:** Anna Marie Hugon, Thaddeus G. Golos

**Author notes:** Correspondence: Thaddeus G. Golos, Ph.D. University of Wisconsin-Madison Wisconsin National Primate Research Center 1223 Capitol Ct. Madison, WI 53715-1299 U.S.A.

## Abstract

Listeria monocytogenes (Lm) is a food-borne pathogen associated with serious pregnancy complications, including miscarriage, stillbirth, preterm birth, neonatal sepsis, and meningitis. Although Lm infection within the gastrointestinal (GI) tract is well studied, little is known about the influence sex hormones may have on listeriosis. Estradiol (E2) and progesterone (P4) not only have receptors within the GI tract but are significantly increased during pregnancy. The presence of these hormones may play a role in susceptibility to listeriosis during pregnancy.

Caco-2 cell monolayers were grown on trans-well inserts in the presence of E2, P4, both E2 and P4, or no hormones (control). Cells were inoculated with Lm for 1 hour, before rinsing with gentamycin and transfer to fresh media. Trans-epithelial resistance was recorded hourly, and bacterial burden of the apical media, intracellular lysates, and basal media were assessed at 6 hours post inoculation.

There were no significant differences in bacterial replication when directly exposed to sex steroids, and Caco-2 cell epithelial barrier function was not impacted during culture with Lm. Addition of P4 significantly reduced intracellular bacterial burden compared to E2 only and no hormone controls. Interestingly, E2 only treatment was associated with significantly increased Lm within the basal compartment, compared to reduction in the intracellular and apical layers.

These data indicate that increased circulating sex hormones alone do not significantly impact intestinal epithelial barrier integrity during listeriosis, but that addition of P4 and E2, alone or in combination, was associated with reduced epithelial cell bacterial burden and apical release of Lm.

**Summary sentence:** Progesterone and estradiol inhibit infection of Caco-2 intestinal epithelial cells by *Listeria monocytogenes*.

## Introduction

A ubiquitous environmental bacterium, *Listeria monocytogenes* (Lm) is primarily acquired via consumption of contaminated food that can cause gastrointestinal illness, meningitis, and sepsis [1]. The elderly, immunocompromised, neonates and pregnant people have increased susceptibility to listeriosis [2]. Gestational listeriosis is associated with serious pregnancy complications, including miscarriage, stillbirth, preterm birth, and neonatal sepsis and meningitis [3–6]. While Lm is known to cause significant fetal morbidity, it often goes unrecognized in the mother until infection at the maternal-fetal interface (MFI) causes adverse pregnancy outcomes (APOs) [7]. Furthermore, the determinants of increased susceptibility to infection with Lm during pregnancy remain undefined.

Within the GI tract, the gram-positive organism relies upon its dense peptidoglycan coating to survive the highly acidic environment of the stomach, until it reaches the intestinal villi where initial infection of the intestinal epithelium occurs [8, 9]. Within the intestinal epithelium, Lm must overcome multiple barriers to infection including a host associated layer of mucus, the layers of epithelial enterocytes, immune cells, cytokines, metabolites, hormones, and other endogenous microorganisms, to achieve infection of the tissues lining the GI tract.

Epithelial tissues infected by Lm contain polarized cells that form a tight barrier to pathogen translocation. Tight junctions are essential for establishing a barrier to restrict the mixing of the two compartments [10]. Within the gut lumen, the intestinal epithelium forms a cellular layer between gastric contents and the circulatory systems of the body. Lm invades host cells through utilization of the bacterial surface proteins internalin A (InlA) and internalin B (InlB) [11]. Upon binding to epithelial cells, InlA and InlB activate signaling pathways of the host surface cell receptors and recruitment of materials needed for endocytosis. The signaling cascade leads to recruitment of actin and actin polymerization. During bacterial entry interactions with E-cadherin favor interaction of the E-cadherin cytoplasmic tail and the actin cytoskeleton, while entry via c-Met stimulates a signaling cascade to promote depolymerization of actin, which completes the process of bacterial internalization. It is worth noting that both of these receptors have been documented in colonic tissues [12, 13]. Within the host cell, expression of ActA aids in polymerization of actin and generation of an actin tail to propel the bacterium through the cell cytoplasm and through membrane protrusions to enter neighboring cells[11]. Spreading within polarized epithelial cell layer is dependent upon internalin C (InlC) which promotes the formation of protrusions through inhibition of cellular Tuba and N-WASP which have been shown to modulate the structure apical junctions[14, 15].

By successfully invading i the GI epithelial barrier through a cycle of replication and shedding into the gut lumen, Lm gains access to the lymphatic system through a process known as transcytosis [8, 11, 16]. Transcytosis is a form of cellular trafficking that Lm utilizes to move from one membrane to another, in this case from the apical layer to the basal layer of the GI epithelium, to ultimately enter the bloodstream [17]. Once in the bloodstream, Lm may circulate until it can access the intervillous space of the placenta, where molecules are transferred from mother to fetus including amino acids, fatty acids, glucose, and oxygen to underpin fetal development [18]. Lm in the intervillous space of the MFI is able to establish severe placental infection which ultimately causes acute inflammation, chorioamnionitis, and necrosis [19].

While the invasion and intracellular phases of Lm at the intestinal epithelial layer have been well described, the determinants of increased susceptibility to listeriosis during pregnancy remain unknown [20–25]. To evaluate these determinants, it is important to consider potential influences of the maternal gut environment on listeriosis. During the course of a normal and healthy pregnancy, there are dramatic changes in the mother’s hormonal, metabolic, and immunological homeostasis. An important component of the pregnant state is elevation in the levels of the sex hormones E2 and P4 [26]. Circulating levels of P4 are elevated during the three trimesters of pregnancy, peaking during the third trimester, while the levels of E2 increase slowly before a rapid increase near the end of gestation [27]. These changes also are thought to alter immune responses during pregnancy [26, 28–30]. During gestation, there is an overall decrease in pro-inflammatory cytokines and increase in counterregulatory cytokines [31]. These immune changes allow tolerance to potentially immunogenic paternal antigens in order to maintain a successful pregnancy. However, the loss in inflammatory signaling during specific stages of pregnancy creates a permissive state in which pathogen invasion does not trigger inflammation and a prompt robust immune response [26].

The modulation of inflammation plays an important role in not only the MFI, but also the GI tract. Researchers have demonstrated that sex steroids and theirs receptors serve an important role in the GI tract and contribute to the progression of a number of GI diseases, including inflammatory bowel disease (IBD), irritable bowel syndrome (IBS) and a variety of GI tract cancers [32–35]. IBD is an incompletely understood intestinal inflammatory disease, and is clinically characterized by a “leaky” GI tract, hypothesized to permit bacterial product translocation and inflammation [36]. Studies suggest that E2 may protect against acute colitis through the activation of ERβ [37, 38]. One study demonstrated that treatment with E2 reduced inflammation in the colon in mice [39]. Research has also shown progesterone to inhibit IBD disease by improving gastrointestinal barrier function during pregnancy [40].

The presence of receptors for these hormones in intestinal epithelial cells has been long known [41–43], however their effects on epithelial barrier function during disease remains unclear. While studies have shown alterations to GI motility during pregnancy, evidence supports a mostly direct effect of P4 on intestinal smooth muscle, rather than progesterone-mediated pathways of gastric motility [33]. P4 has been documented to increase transepithelial electrical resistance in primary human colon tissues and Caco-2 cells through upregulating expression of the tight junction protein occludin [44]. In addition, E2 has been shown to decrease permeability through modulation of paracellular permeability and tight junctions [45]. However, whether sex steroids affect the transcellular pathway in the intestinal epithelium remains undefined.

It is possible that circulating sex hormones impact the maternal gut microenvironment susceptibility to infection with Lm through incompletely defined mechanisms. We hypothesized that the presence of P4 alone would increase TEER, while E2 alone would decrease TEER. We also hypothesized that treatment with both P4 plus E2 would have decreased Lm within the intracellular and basal compartments. Receptors for both hormones have been documented in the human epithelial colorectal cancer (Caco-2) cell line making it an ideal model for examining the direct effects of P4 and E2 on Lm replication within the GI tract [32, 41, 44, 46]. Our study aimed to build upon the existing knowledge by evaluating the impact of E2 and P4 during Lm infection of intestinal epithelial monolayers using a Caco-2 model.

## Methods

### Cell Culture

Human Caco-2 cells (HTB37; American Type Culture Collection) purchased from ATCC (Manassas, VA, USA) were maintained in Dulbecco’s Modified Eagle’s medium (DMEM, phenol red-free, Thermo Fisher Scientific, Waltham, MA, USA) supplemented with 20% charcoal-stripped FBS, 2 mM L-glutamine, 10 mM HEPES, 100 unit/mL penicillin and 100 µg/mL streptomycin (Thermo Fisher Scientific, Waltham, MA, USA) at 37°C under room air/5 % CO_2_ in a humidified incubator. Cells were grown in 25 cm² culture flasks (Corning, Corning, NY, USA). The medium was changed every three days. The cells were harvested for passage or plating with 0.25% trypsin-EDTA (Thermo Fisher Scientific, Waltham, MA, USA).

To initiate infection experiments, the cells were harvested from confluent cell cultures and suspended in DMEM containing 10% fetal bovine serum and 1% non-essential amino acids. 24-well tissue culture plates containing 8 μm pore Polyethylene (PET) membrane inserts (catalogue number: 25-289, Corning, Corning City, NY, USA) were seeded with 3.5 × 10^4^ cells per well and cultured to confluence with a final density of approximately 4 × 10^5^ cells per well. Cells were then differentiated into a monolayer using Corning BioCoat Intestinal Epithelial Environment and protocol (catalog number: 355057, Corning, Corning City, NY, USA).

Hormone stocks of E2 or P4 were dissolved in ethanol to a stock concentration of 1 mg/mL, with further dilutions made in Modified Eagle’s medium (DMEM, phenol red-free, Thermo Fisher Scientific, Waltham, MA, USA) supplemented with 20% charcoal-stripped FBS, 2 mM L-glutamine, and 10 mM HEPES with no antibiotics. Wells were then treated with 2.50 ng/ml E2, 40 ng/ml P4, both, or no hormones as a control. Hormone concentrations were based upon circulating concentrations during pregnancy as reported in humans [47]. Since both the P4 and E2 were dissolved in absolute ethanol then diluted into media, an appropriate amount of ethanol was added to control wells. No impact of ethanol addition alone (<0.1%) was noted in preliminary studies. Cells were incubated in hormones for 24 hours prior to experimentation.

### Bacterial Culture

*L. monocytogenes* 2203S (wild type [WT]; serovar 4b [48]) was cultured overnight at 37◦C in Tryptic Soy Broth (TSB) (Becton Dickinson, Sparks, MD). Bacterial concentration was estimated using optical density measurements at a wavelength of 595 nm. Fresh bacterial cultures were washed and resuspended in Dulbecco’s modified Eagle medium (DMEM, phenol red-free, Thermo Fisher Scientific, Waltham, MA, USA) containing 10% fetal bovine serum (D10F) before addition to Caco-2 monolayers at a multiplicity of infection (MOI) of 1. The infectious dose was confirmed retrospectively by culturing tenfold serial dilutions of the inoculum in PBS on blood agar plates. Cells were incubated with Lm for 1 hour, then media were removed and replaced by D10F containing 50 μg/mL gentamicin (Thermo Fisher Scientific, Waltham, MA, USA) to kill extracellular bacterial cells (Fig 1). Gentamycin media were then removed, cells were washed, and transepithelial electrical resistance and infection studies were initiated.

**fig 1.**
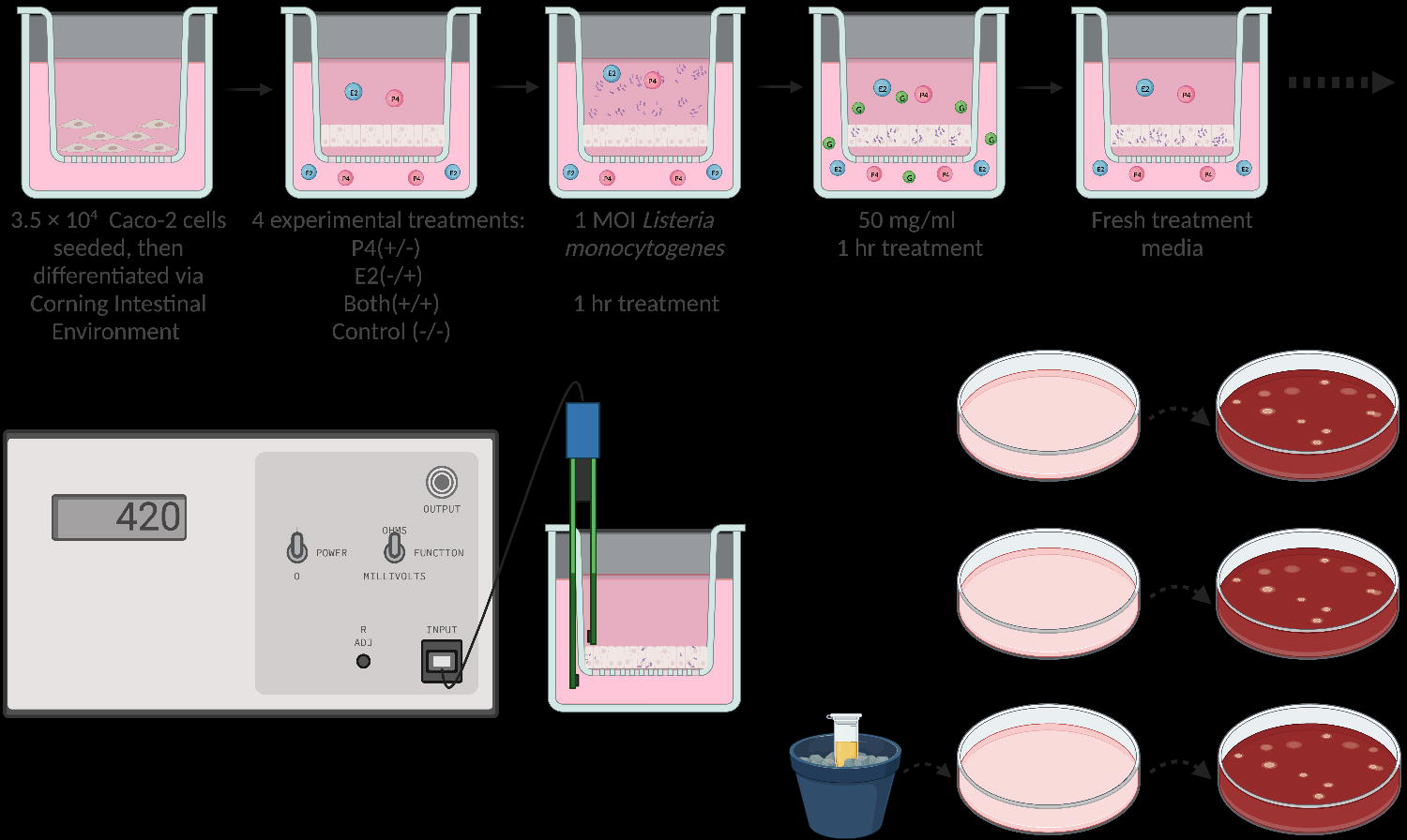

### Barrier Function

Barrier integrity was assessed using Trans-Epithelial Electrical Resistance (TEER) of polarized monolayers measured using the Millicell-Electrical Resistance System (ERS, Millipore, Billerica, MA, USA) and those with a minimum TEER of 200 Ο/cm^2^ (range, 200 to 600 Ο/cm^2^) were used for translocation experiments.

Pilot experiments were conducted to determine the appropriate length of time for monitoring bacterial translocation through Caco-2 monolayers, and hourly timepoints for 6 hours following exposure were chosen to capture the full progression of replication and translocation. Inserts with no cells as well as seeded inserts with no bacteria were used as negative controls for TEER. Each hormone treatment group was performed in duplicate wells and the experiment was replicated three times.

At 6 hours post exposure, the apical and basal media were separately collected, serially diluted, and plated on blood agar for the enumeration of Lm following 24 and 48 hours. The cells were resuspended in media and mechanically disrupted using the freeze-thaw method to release intracellular bacteria [49]. Lysates were serially diluted in PBS and plated on blood agar (Fig 1).

### Statistical analysis

All data analysis and graphs were prepared using Prism 9 software (Graph-Pad Software Inc., San Diego, CA, USA). Data from repeated experiments are presented as mean and standard error (SEM). Where appropriate, Tukey’s multiple comparison test was used to identify statistically significant differences (P<0.05).

## Results

### Bacterial Replication

First, we cultured Lm in TSB Media with 2.5 ng/ml E2, 40 ng/ml P4, both hormones, or no hormones (serving as control) to determine the impact of each treatment on survival and replication of Lm. 10^4^ CFU Lm were added followed by incubation at 37 °C. Bacterial levels (CFU/ml) of Lm was assessed using OD595 hourly for 6 hours following addition of Lm to hormone treated media. ANOVA with Turkey’s multiple comparison test found no significant differences in bacterial levels between hormonal treatment groups during 6h of culture (Fig 2).

**fig 2.**
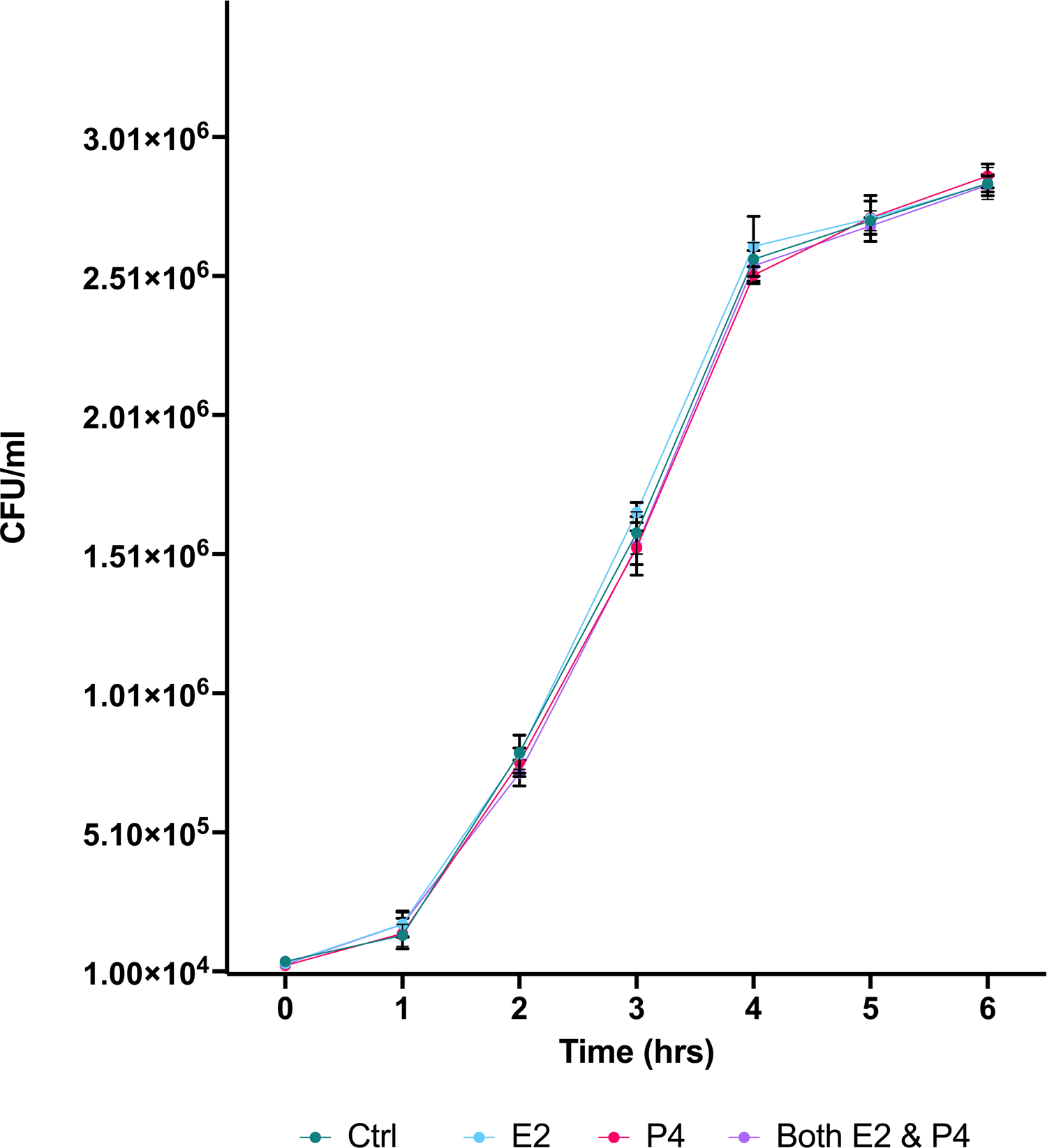

### Barrier Function

We then examined the impact of hormone treatment on intracellular replication and barrier function with transcytosis of Lm by culturing cells on Transwell filter inserts on which Caco-2 cells were cultured to confluence. Culture plates were treated with media containing E2, P4, both, or neither hormone. Following 24 hours, 10^5^ CFU (1 MOI) of bacteria were added to the apical wells. The CFU of the inoculum was confirmed using culture-based methods. Plates were incubated for 1 hour, and then washed with medium containing 50 μg/mL gentamicin to kill extracellular Lm. Filter inserts were then transferred into 24-well tissue culture plates containing fresh media with the same hormones as prior to inoculation. Trans-epithelial resistance (TEER) was recorded hourly at 1 - 6 hours post-exposure (Fig 3). 2-way ANOVA and Tukey’s multiple comparisons test of each treatment group compared with controls (no hormones) revealed no significant changes in barrier function with hormone treatment.

**fig 3.**
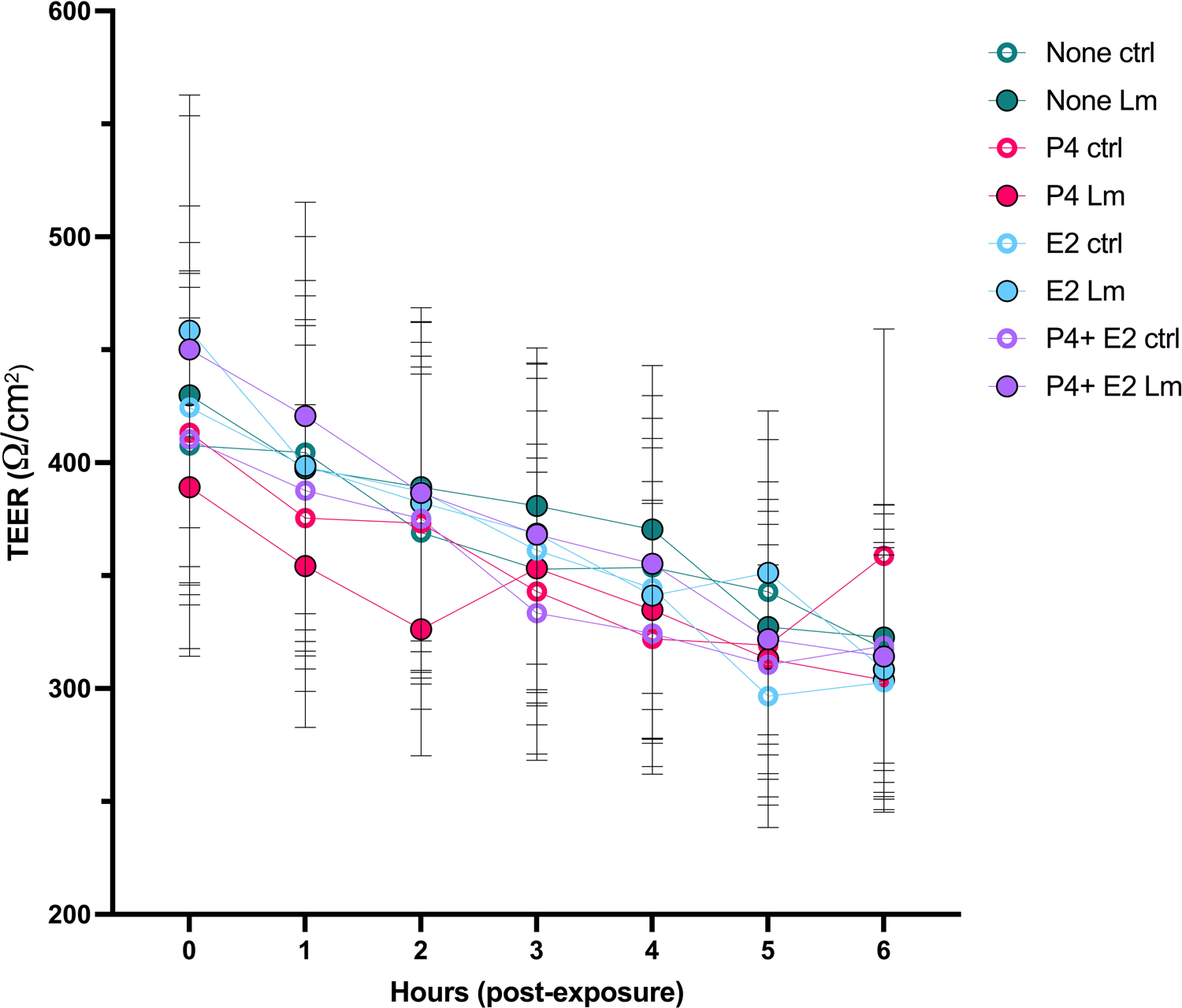

### Bacterial burden by Compartment

Following 6 hours incubation, the apical and basal compartment media were collected and plated to determine bacterial burden, the inserts were removed, and epithelial cells were lysed to determine intracellular bacterial burden (Fig 4). In the apical media, there was significantly less bacterial burden with P4 treatment only, compared to E2 only (p = 0.0014), E2 plus P4 (p = 0.0004), or no hormone controls (p = <0.0001). Both the E2 only (p = 0.0034) and both hormone treatment groups (p = 0.0112) also had significantly lower bacterial burden compared to no hormone controls.

**fig 4.**
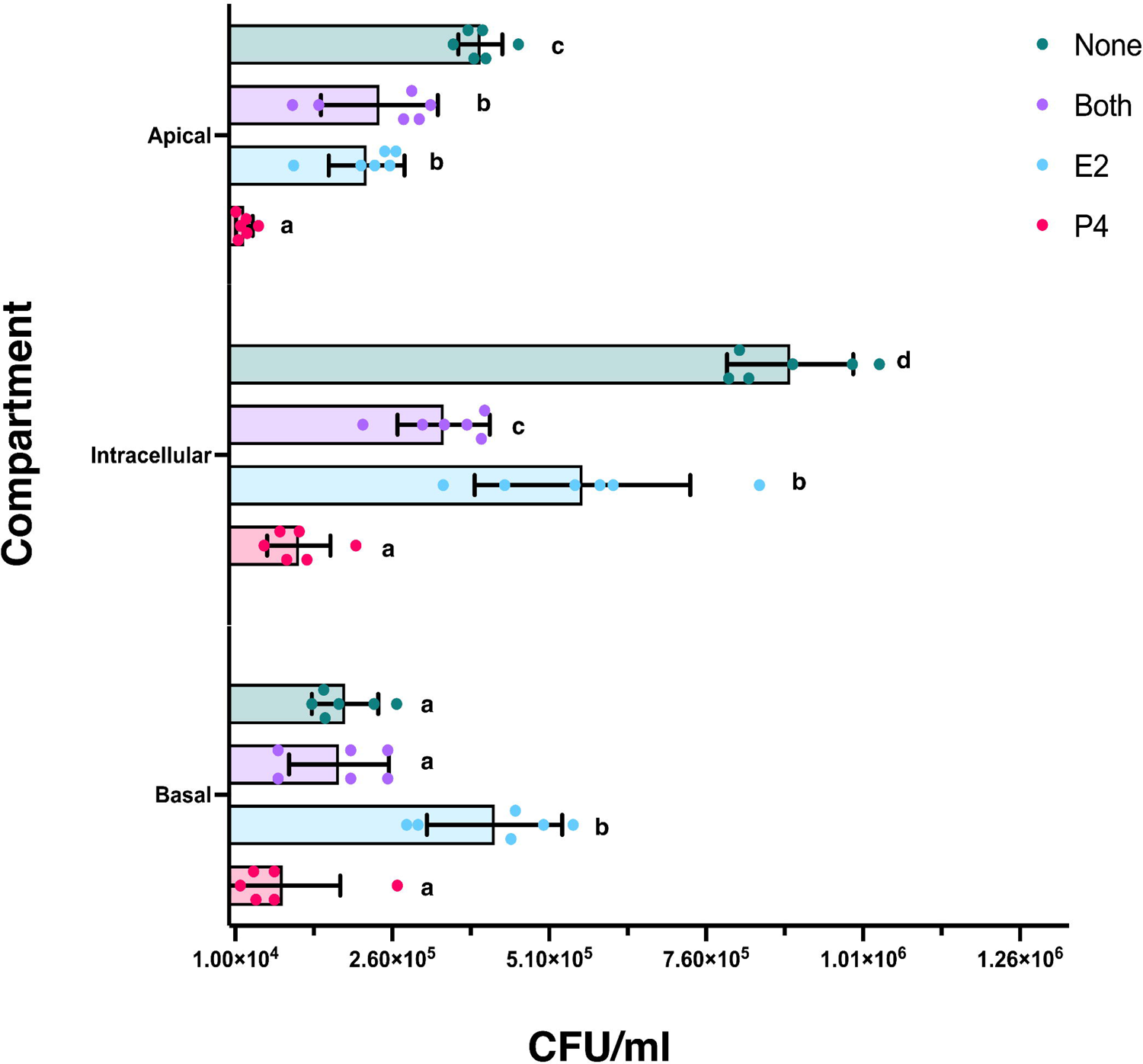

The intracellular lysate had similar results, with significantly lower bacterial burden with P4 treatment only, compared to E2 only (p = <0.0001), E2 plus P4 (p = 0.0001), or no hormone controls (p = <0.0001). Within the lysates, the E2 only treatment group also had significantly more Lm compared to E2 plus P4 (p = 0.0002), but significantly less intracellular bacteria than the no hormone controls (p = <0.0001); cells treated with both hormones also displayed significantly lower bacterial burden than controls (p = <0.0001).

Within the basal media, the P4 only treatment group had significantly less bacteria compared to the E2 only treated group (p = <0.0001) and was lower than the E2 plus P4 or no hormones controls, although that difference did not achieve significance. Furthermore, the E2 only treated basal media had significantly increased bacterial burden when compared to either P4 only, E2 plus P4, or no hormone controls (p = <0.0001).

## Discussion

### Sex Steroids and Bacterial Replication

In this study, we tested the ability of Lm to infect, and alter the barrier resistance of intestinal epithelial cell monolayers in the presence of P4 and E2 to elucidate the potential impact of pregnancy sex steroids on passage of Lm across the epithelium of the GI tract. While the intracellular phase of Lm infection has been extensively studied, little is known about the impact of sex steroids during infection of Lm within the GI tract. To rule out any effect of sex hormones directly on Lm replication, the impact of P4 and E2 on bacterial growth was tirst assessed in TBS culture. The data indicated that the replication of Lm is not directly affected by these hormones (Fig 2), contirming that sex steroids used in subsequent culture experiments do not directly impact Lm replication.

### Sex Steroids and Barrier Function

To asses barrier function, we utilized a cell line derived from a colon adenocarcinoma (Caco-2), which differentiates into enterocyte-like cells under specific culture conditions and is a widely used model to assess intestinal epithelial permeability [50]. These cells develop into a polarized monolayer which form a border between the apical and basal compartments. Polarized epithelial cells are characterized by their ability to conduct endocytosis at either the apical or basal membranes [51]. The cell monolayer is organized similar to a honeycomb pattern, with junctions along all sides of the cells [50]. Tight junctions form the border at the basolateral cell surface domains in polarized epithelia, and support the maintenance of cell polarity by restricting intermixing of apical and basolateral transmembrane components [10]. Tight junctions include occludins and claudins, forming the zonula occludens [52]. These junctions modulate permeability between the intestinal epithelial cells, as well as intersect with signaling mechanisms that direct epithelial-cell polarization and the formation of apical and basal domains that are morphologically and functionally distinct [10]. Furthermore, the basal extracellular matrix constructed by epithelial cells in coordination with their junctions creates a barrier of separation between GI contents and circulatory networks of the body and is a barrier to bacterial dissemination into the bloodstream. This epithelial layer separates the contents of the intestinal lumen microenvironment of the GI tract including the GI contents, host-associated mucus, and microbial presence from the underlying circulatory system and lymphatic network of the intestinal villi. Each of these compartments has unique properties and represents various stages of Lm replication and release. The epithelial layer is where Lm replicates, and upon release of Lm from the cells the bacterium can freely reenter the lumen via the apical surface or escape the basolateral membrane to access the host circulation.

In our model, Lm readily infected Caco-2 cells and multiplied intracellularly but did not significantly decrease TEER of the Caco-2 monolayer as compared to uninfected cells. While there was no clear impact of sex steroids or Lm on transepithelial resistance, we found that within the apical and intracellular compartments, treatment with E2 or P4 independently led to a decrease in Lm burden. This effect of P4 was confirmed in the basal layer, with decreased Lm burden with P4 treatment, however treatment with E2 led to significantly increased Lm within the basal compartment, suggesting a more complex action of hormones on Lm passage across the epithelial barrier.

### Sex Steroids and Barrier Function

Through use of an intestinal epithelium cell culture model, we further found that the presence of P4 and/or E2 had no significant impact by themselves on transepithelial barrier resistance during the experimental period (Fig 3). These data indicate that the presence of sex steroids alone does not significantly impact epithelial barrier resistance during Lm infection of the GI tract, although limitations of this culture system include that the epithelial environment in the gut *in vivo* is much more complex, including exposure to hormones over months, and that the adenocarcinoma cells may not accurately recapitulate the *in vivo* enterocytes.

Other studies examining the effects of P4 and E2 on Caco-2 barrier function found conflicting results. Salomon et. al examining Caco-2 cell brush-border membranes with P4 or E2 treatment found no effect on TEER of hormone treatment [46]. That study examined TEER function 5-, 10-, and 30-days following hormone treatment with P4 (310ng/mL) or E2 (270 ng/mL), with no differences in barrier function occurring at each of the time points. However, Zhou et al treated Caco-2 cells with 20 and 125 ng/mL of P4 for 24 h, and those authors found that TEER values were significantly increased following treatment with both P4 doses [44]. These data suggest that a longer treatment period is not required, but that increased concentrations of P4 may promote changes to barrier permeability. While we selected the dose of hormones in our experiments based on circulating levels in human pregnancy, future studies could examine the impact of other hormone doses prior to exposure to Lm.

One potential mechanism through which sex steroids could indirectly impact permeability is through modulation of cytokine signaling and inflammation. In the aforementioned study, the authors also examined serum from pregnant women during early gestation, late gestation, and post-partum stages and found that progesterone with pro-inflammatory cytokine levels and showed that TNF-α, IL-6 and IL-1β were significantly reduced during the third trimester compared to postpartum [44]. These results indicate that the production of pro-inflammatory cytokines might be impacted through an indirect progesterone-mediated mechanism. A progesterone-mediated inhibition of pro-inflammatory cytokines could lead to decreased inflammation, diminished pathogen recognition, and immune cell activation, thus dampening the host response to infection. The impact of pro-inflammatory cytokines extends beyond immunologic activities and have been shown to affect epithelial barrier function [36, 53, 54], a key restraint on dissemination of Lm across the GI tract. Increased levels of P4 during pregnancy may prevent dissemination of Lm or reduce systemic inflammation through currently undefined mechanisms. We may not see an impact of hormones on TEER in our study as the Caco-2 cells are a limited model of the GI tract, i.e., only the intestinal epithelium. This does not account for changes to circulating cytokines as well as many other factors, including the complexity of the intestinal mucosa.

### Sex Steroids and Bacterial Burden

Through quantification of Lm within the distinct basal, intracellular, and apical compartments of the Transwell culture system, we found a complex and significant impact of sex steroids on Lm replication and burden (Fig 4). In the apical media, addition of E2 or P4 independently led to a significantly decreased burden of Lm compared to controls. The combination of both hormones decreased Lm replication and release into the apical layer compared to controls, but the combination of E2 and P4 was not different from E2 alone, although significantly greater than P4 alone.

With intracellular lysates, either P4 or E2 treatment alone led to significantly reduced cellular Lm burden compared to controls. Treatment with both hormones also led to a significantly greater decrease in intracellular Lm compared to E2 only. This suggests that inhibition of Lm replication by P4 was additive to inhibition by E2, although the effect of P4 in the absence of E2 was also significantly greater than the effect of E2 plus P4.

The impact of sex steroids on Lm in the basal compartment was different from that seen in the apical or intracellular compartments. Lm burden in cells treated with E2 alone was statistically significantly higher than the other treatment groups. Lm burden in the P4 only group was similar to the apical levels, however the relatively lower control or E2 plus P4 groups resulted in no significant effect of P4 on Lm levels. This implies that there is differential release of Lm from the basal and the apical surfaces of the Caco-2 cells in this transwell chamber system. Collectively, across the apical, intracellular and basal compartments, these data confirm that P4 decreases Lm intracellular replication which is reflected by reduced release of Lm into the apical layer and reduced bacterial burden in the intracellular compartment.

In considering why the release of Lm into the basal compartment differs from the apical or intracellular compartments, one needs to consider Lm movement between and within epithelial cells. Motility of Lm within the epithelial layer is dependent on the host cell actin cytoskeleton and access to surface proteins such as E-cadherin and c-Met, which are normally on the basal side of intestinal cells [11]. Lm has been shown take advantage of apoptoic extrusion, which is a mechanism to remove dying or unwanted cells from an epithelium layer while preserving the barrier function [55]. Lm takes advantage of the temporarily exposed basal surface binding proteins to gain entry into the cell. Lm virulence factors such as InlC have been shown to further promote formation of cell protrusions, which aids in bacterial replication, comet tail assembly, and disrupts the structure of apical junctions in epithelial cells [15]. It is possible that E2 supports increased release of Lm into the basal compartment by indirectly altering and exposing tight junctions by increasing epithelial cell renewal and junction remodeling. E2 has been shown to have a protective effect on the gut epithelium, reducing inflammation through activation of Estrogen receptor-β (ERβ) [37–39]. ERβ expression in the GI tract has been reported to be higher in females compared to males [42]. Further, ERβ signaling has been shown to modulate epithelial barrier function [38, 40, 45]. One study demonstrated a reduction in ERβ mRNA expression and an increase in gut permeability prior to the onset of colitis in two animal models of spontaneous colitis [38]. The authors also used RT-PCR and electric cell-substrate impedance sensing of HT-29 and T84 colonic epithelial monolayers and found increased barrier resistance with E2 treatment. These data suggest that not only is more advanced modeling required, but that alterations to intestinal permeability are dependent on signaling pathways which are indirectly affected by the presence of sex steroids and their receptors.

These data contribute to the beginning of our understanding of the determinants of susceptibility to listeriosis during pregnancy, as they examine the direct effect of sex steroids on Lm infection of the intestinal epithelial barrier and dissemination of Lm throughout the body. However, there are multiple elements to the barrier between the intestinal lumen and the host blood space. The basal extracellular matrix produced by epithelial cells and their lateral intercellular junctions creates a barrier of separation between GI contents and circulatory networks of the body. In our studies, the presence of Lm within the basal layer represents successful intracellular replication and transcytosis across the intestinal epithelial monolayer.

## Conclusions

In summary, we have demonstrated that Caco-2 cells are readily infected with *L. monocytogenes* in vitro and that infected monolayers do not exhibit decreased monolayer integrity, as measured by TEER. In addition, there were no significant differences on the epithelial barrier function during listeriosis with exposure to sex steroids. However, the data indicate that treatment with either E2 or P4 generally decreases Lm replication and release, although there may be differential steroid hormone regulation release of Lm into the apical and basal compartments.

These data suggest that any impact of elevated circulating sex hormones in pregnancy on increased susceptibility to listeriosis does not rely solely on their impact on intestinal epithelial cell barrier function. Indeed, sex steroids actually appear to inhibit Lm replication in intestinal epithelial cells. Our study expands upon our understanding of the complexities of Lm infection and susceptibility of the maternal GI tract. It is well known that pregnancy is a risk factor for listeriosis, and that listeriosis in pregnancy is associated with a spectrum of adverse pregnancy outcomes. We have also recently confirmed susceptibility to listeriosis in a pregnant macaque model, and that pregnancy also impacts susceptibility to gut dysbiosis in gestation [56]. It is possible that the maternal gut microenvironment may play a role in dispersion of Lm outside of the intestinal tract, with commensal microbes influencing Lm survival and invasion of epithelial tissue. Further studies are needed into the possibility that sex steroid-induced changes in the intestinal microbiome during pregnancy, or hormonal impact on other elements of the intestinal wall or immune system, may be involved on conferring increased susceptibility to listeriosis.

## Acknowledgements

We thank Dr. Chuck Czuprynski, UW-Madison Dept. of Pathobiological Sciences for helpful discussion on experimental design, the Dr. Federico Rey Laboratory in the UW-Madison Dept. of Pathobiological Sciences for use of their equipment for determining TEER, and we thank Dr. Sophia Kathariou of North Carolina State University for generous donation of clinical strain LM2203S.

## Contributions

TGG, and AMH designed the study. AMH conducted experiments and analyzed the data. AMH and TGG drafted the manuscript. AMH created all figures and tables. All authors read and approved the final manuscript.

## Disclaimer

The contents of this manuscript are solely the responsibility of the authors and do not represent the official views of the NIH.

## Data Availability

The authors confirm all data supporting these findings are available within the manuscript or cited materials.

## Supplemental Data

**Supplemental Table 1** Statistical Analyses

