## Supplementary material for "*Listeria monocytogenes* infection in intestinal epithelial Caco-2 cells with exposure to progesterone and estradiol-17beta in a gestational infection model": Figure & Table Legends

**Figure 1** Image depicting the experimental timeline of each well. Each well features a transmembrane insert and media indicated by pink. Sex hormones are depicted by colored spheres, with E2 and P4 colored blue and pink, respectively. Gentamycin is represented by smaller green spheres, and Lm is denoted by purple rods. Following exposure, gentamycin, and washing, TEER was measured hourly for 6 hours then followed by plating of the apical media, basal media, and cell layer lysate on blood agar plates for quantification at 24 and 48 hours.

**Figure 2** The growth of Lm (CFU/ml) during 6-hour incubation following exposure to E2, P4, both E2 and P4, or no hormone (control). The mean +/- standard error of the mean (SEM) is indicated.

**Figure 3** The graph depicts the average TEER values of cells grown on 8 $\mu$m pore inserts for 6 hours post-exposure to Lm. The treatment groups are color coded and Lm exposure is indicated by a filled symbol. The mean +/- SEM is presented.

**Figure 4** The scatterplot depicts the quantity of Lm (CFU/ml) in the apical layer, intracellular lysates, and basal layer. The treatment groups are color coded. The height of the bar indicates the mean of each treatment group, and the bars indicate standard error of the mean (SEM). Values that do not share superscripts are significantly different.

**Supplemental Table 1** Statistical Analysis
