## Supplemental Data Table 1 for "*Listeria monocytogenes* infection in intestinal epithelial Caco-2 cells with exposure to progesterone and estradiol-17beta in a gestational infection model"

| Tukey's multiple co | Mean Diff. | 95.00% CI ( | Below thres | Summary | Adjusted P Value | |
|---|---|---|---|---|---|---|
| Apical | | | | | | |
| P4 vs. E2 | -194683 | -327094 to | Yes | ** | 0.0014 | abc |
| P4 vs. Both | -215017 | -347427 to | Yes | *** | 0.0004 | abc |
| P4 vs. None | -375850 | -508260 to | Yes | **** | <0.0001 | abc |
| E2 vs. Both | -20333 | -152744 to | No | ns | 0.9772 | |
| E2 vs. None | -181167 | -313577 to | Yes | ** | 0.0034 | ac |
| Both vs. None | -160833 | -293244 to | Yes | * | 0.0112 | abc |
| Intracellular | | | | | | |
| P4 vs. E2 | -452000 | -584410 to | Yes | **** | <0.0001 | abcd |
| P4 vs. Both | -230833 | -363244 to | Yes | *** | 0.0001 | abcd |
| P4 vs. None | -782833 | -915244 to | Yes | **** | <0.0001 | abcd |
| E2 vs. Both | 221167 | 88756 to 35 | Yes | *** | 0.0002 | abc |
| E2 vs. None | -330833 | -463244 to | Yes | **** | <0.0001 | abc |
| Both vs. None | -552000 | -684410 to | Yes | **** | <0.0001 | ab |
| Basal | | | | | | |
| P4 vs. E2 | -337667 | -470077 to | Yes | **** | <0.0001 | cd |
| P4 vs. Both | -89667 | -222077 to | No | ns | 0.2884 | |
| P4 vs. None | -99500 | -231910 to | No | ns | 0.2048 | |
| E2 vs. Both | 248000 | 115590 to 3 | Yes | **** | <0.0001 | abc |
| E2 vs. None | 238167 | 105756 to 3 | Yes | **** | <0.0001 | abc |
| Both vs. None | -9833 | -142244 to | No | ns | 0.9973 | |

| Test details | Mean 1 | Mean 2 | Mean Diff. | SE of diff. | N1 | N2 | q | DF |
|---|---|---|---|---|---|---|---|---|
| Apical | | | | | | | | |
| P4 vs. E2 | 24483 | 219167 | -194683 | 50108 | 6 | 6 | 5.495 | 60 |
| P4 vs. Both | 24483 | 239500 | -215017 | 50108 | 6 | 6 | 6.069 | 60 |
| P4 vs. None | 24483 | 400333 | -375850 | 50108 | 6 | 6 | 10.61 | 60 |

| | | | | | | | | |
|---|---|---|---|---|---|---|---|---|
| E2 vs. Both | 219167 | 239500 | -20333 | 50108 | 6 | 6 | 0.5739 | 60 |
| E2 vs. None | 219167 | 400333 | -181167 | 50108 | 6 | 6 | 5.113 | 60 |
| Both vs. None | 239500 | 400333 | -160833 | 50108 | 6 | 6 | 4.539 | 60 |
| Intracellular | | | | | | | | |
| P4 vs. E2 | 111167 | 563167 | -452000 | 50108 | 6 | 6 | 12.76 | 60 |
| P4 vs. Both | 111167 | 342000 | -230833 | 50108 | 6 | 6 | 6.515 | 60 |
| P4 vs. None | 111167 | 894000 | -782833 | 50108 | 6 | 6 | 22.09 | 60 |
| E2 vs. Both | 563167 | 342000 | 221167 | 50108 | 6 | 6 | 6.242 | 60 |
| E2 vs. None | 563167 | 894000 | -330833 | 50108 | 6 | 6 | 9.337 | 60 |
| Both vs. None | 342000 | 894000 | -552000 | 50108 | 6 | 6 | 15.58 | 60 |
| Basal | | | | | | | | |
| P4 vs. E2 | 85333 | 423000 | -337667 | 50108 | 6 | 6 | 9.53 | 60 |
| P4 vs. Both | 85333 | 175000 | -89667 | 50108 | 6 | 6 | 2.531 | 60 |
| P4 vs. None | 85333 | 184833 | -99500 | 50108 | 6 | 6 | 2.808 | 60 |
| E2 vs. Both | 423000 | 175000 | 248000 | 50108 | 6 | 6 | 6.999 | 60 |
| E2 vs. None | 423000 | 184833 | 238167 | 50108 | 6 | 6 | 6.722 | 60 |
| Both vs. None | 175000 | 184833 | -9833 | 50108 | 6 | 6 | 0.2775 | 60 |
